# Functional Somatotopy of Lumbar Dorsal Rootlets and its Role in Selective Recruitment via Lateral Spinal Cord Stimulation

**DOI:** 10.64898/2026.06.18.733242

**Authors:** M Del Brocco, GJ Ansah, M Duran, S Bhowmick, C Gopinath, MK Jantz, R Bose, SF Lempka, LE Fisher

## Abstract

**Objective:** Lateral spinal cord stimulation (LSCS) is a promising approach for restoring somatosensory feedback in lower-limb amputees, but its spatial selectivity remains limited. Percepts often spread to unintended regions of the residual limb, and reducing electrode contact size may not improve focality. This study investigated whether the anatomical organization of lumbar dorsal rootlets (DR) imposes fundamental constraints on LSCS selectivity.

**Approach:** Acute neurophysiology experiments were performed in six adult cats. Both LSCS and individual DR stimulation were conducted in the same animals. For DR stimulation, bipolar hook electrodes were used to stimulate individual DR, while antidromic compound action potentials (CAPs) were recorded from femoral and sciatic nerve branches instrumented with nerve cuffs. For LSCS, custom 32-contact epidural paddle electrodes were placed over the lateral surface of the spinal cord at corresponding vertebral levels. Recruitment thresholds, dynamic ranges, and response patterns were analyzed across spinal levels, and DR recruitment patterns were directly compared to those evoked by LSCS within the same animals.

**Main results:** A clear rostrocaudal organization was observed across spinal levels during stimulation of individual DR, with femoral branches predominantly recruited at L4–L5 and sciatic branches at L6–L7. However, no somatotopic organization was found across DR within each spinal level; individual DR frequently co-activated multiple branches within the same group, and selective recruitment could only be maintained over a narrow dynamic range (median ∼10 µA). LSCS exhibited even a narrower dynamic range (∼5 µA) but closely mirrored DR recruitment patterns, indicating that LSCS activates sensory afferents in a manner determined by the organizational structure of the DR.

**Significance:** These findings demonstrate that the limited spatial selectivity of LSCS can largely be attributed to the coarse organization of DR within each root level rather than due to limitations of epidural electrode design. Moving electrodes intradurally or reducing contact size further is unlikely to substantially improve focality. Instead, improving paddle stability to ensure consistent placement over the appropriate spinal levels may be a more effective strategy for enhancing percept localization.

## INTRODUCTION

Every year, thousands of individuals undergo lower-limb amputation [1], with many experiencing persistent mobility challenges and postural instability while standing and walking with a prosthesis. These impairments are strongly linked to the loss of somatosensory feedback from the missing limb [2]. Tactile inputs from the foot and leg play a critical role in balance, gait, and motor control [3]. Their absence creates a mismatch between intended movement and expected sensory input, degrading prosthesis control and user confidence, and many of the actions performed with the legs are governed by sensory input onto neural circuits in the spinal cord and brainstem. As a result, restoring somatosensory feedback has become a central goal in the development of next generation neuroprosthetic systems for amputees.

Peripheral nerve stimulation has shown promise in evoking sensations perceived in the missing limb and improving prosthesis use [4], [5], [6]. Techniques such as flat-interface nerve electrodes [4], [7], [8], intrafascicular arrays [5], [6], [9], and targeted sensory reinnervation [10] have demonstrated restored tactile and proprioceptive sensations. However, these approaches require complex surgical procedures and may be unsuitable for individuals with peripheral neuropathies or high-level amputations, where distal nerves may be damaged or absent.

An alternative is to stimulate more proximal structures in the nervous system. Our lab has recently demonstrated that spinal cord stimulation, a widely used clinical technique for pain management [11], [12], may be a viable approach for restoring sensation. In particular, lateral spinal cord stimulation (LSCS) has been shown to evoke sensations localized to the distal limbs, including hands and feet, by activating sensory afferents in the dorsal roots [13], [14]. However, these evoked sensations are often diffuse or include off-target percepts in the residual limb [13], [14], suggesting there may be limitations in the spatial selectivity of LSCS. Although current SCS devices use relatively large electrode contacts (area: 6–12 mm^2^), our recent study in cats demonstrated that reducing contact area from ∼0.79 mm^2^ (1.0 mm diameter) to ∼0.20 mm^2^ (0.5 mm) and ∼0.018 mm^2^ (0.15 mm) did not significantly improve selectivity [15]. These findings suggest that anatomical factors, rather than contact size alone, may impose fundamental constraints on focal activation in LSCS. One potential source of these limitations lies in the organization of the dorsal rootlets (DR), which are the primary neural structures activated during LSCS [16].

Beneath the dura of the spinal cord, dorsal roots diverge into smaller structures known as DR before entering the spinal cord [17]. These DR are densely packed and anatomically aligned along the lateral surface of the cord, making them a promising target for selective stimulation [17], [18]. Yet the degree of somatotopic organization across and within DR remains poorly understood. If adjacent DR innervate focal adjacent limb regions (e.g., different toes), even coarse stimulation might suffice for focality. But if neighboring DR project to widely spaced areas (e.g., big toe vs. heel) in an unpredictable or disorganized pattern, then high-precision stimulation would be required to avoid unwanted sensations.

Using neurophysiology experiments, here we investigated whether the functional organization of the lumbar DR could explain the limitations of LSCS focality. Specifically, we compared the nerve recruitment patterns elicited by LSCS versus stimulation of the lumbar DR. In acute experiments in cats, we stimulated individual DR and recorded antidromic compound action potentials (CAPs) from femoral and sciatic nerve branches to characterize rootlet-level recruitment patterns. We then compared these patterns to those evoked by epidural LSCS at matched vertebral levels. This comparison allowed us to assess how epidural stimulation interacts with the underlying rootlet organization, and to what extent LSCS engages similar afferent pathways. Our findings provide new insight into the anatomical basis of sensory recruitment in LSCS and support future efforts to refine stimulation strategies for somatosensory neuroprosthetics.

## METHODS

### Animal preparation

Experiments were conducted in acute terminal preparations in adult cats (n = 6), under the approval of the University of Pittsburgh Institutional Animal Care and Use Committee (IACUC). The same animals were used in our companion study [15]. Anesthesia was induced with an intramuscular injection of ketamine/acepromazine and maintained with 1–2% isoflurane. A tracheotomy was performed to facilitate mechanical respiration, and a catheter was inserted into the carotid artery to enable continuous blood pressure monitoring. Throughout the experiment, physiological parameters including end-tidal CO_2_, oxygen saturation (SpO_2_), heart rate, core body temperature, and blood pressure were monitored and maintained within normal ranges. The animal was placed in a stereotactic frame to ensure surgical stability. Upon completion of data collection, animals were euthanized with an IV injection of Euthasol.

### Peripheral nerve instrumentation

The left femoral and sciatic nerves were exposed, and cuff electrodes (Cortec GmbH, Germany) were placed around their main trunks and accessible branches (Figure 1a, 1d, 1e) to record electroneurogram (ENG) signals. Cuff electrodes were selected with internal diameters of 1, 1.5, 2, 2.5, or 3 mm depending on the size of the nerve branch. Twelve nerves were typically instrumented in each experiment, including the femoral trunk, sciatic trunk, tibial, distal tibial, common peroneal, sural, and saphenous nerves, as well as motor branches innervating the medial and lateral gastrocnemius, biceps femoris, rectus femoris, and vasti muscles. In Cat A, the nerve branch innervating the biceps femoris was not instrumented due to difficulty locating it, and data from the medial and lateral gastrocnemius nerves were excluded from analysis due to excessive noise. A summary of the nerves instrumented is shown in Figure 1a. All cuff electrodes were used to record CAPs evoked by LSCS and DR stimulation.

**Figure 1:**
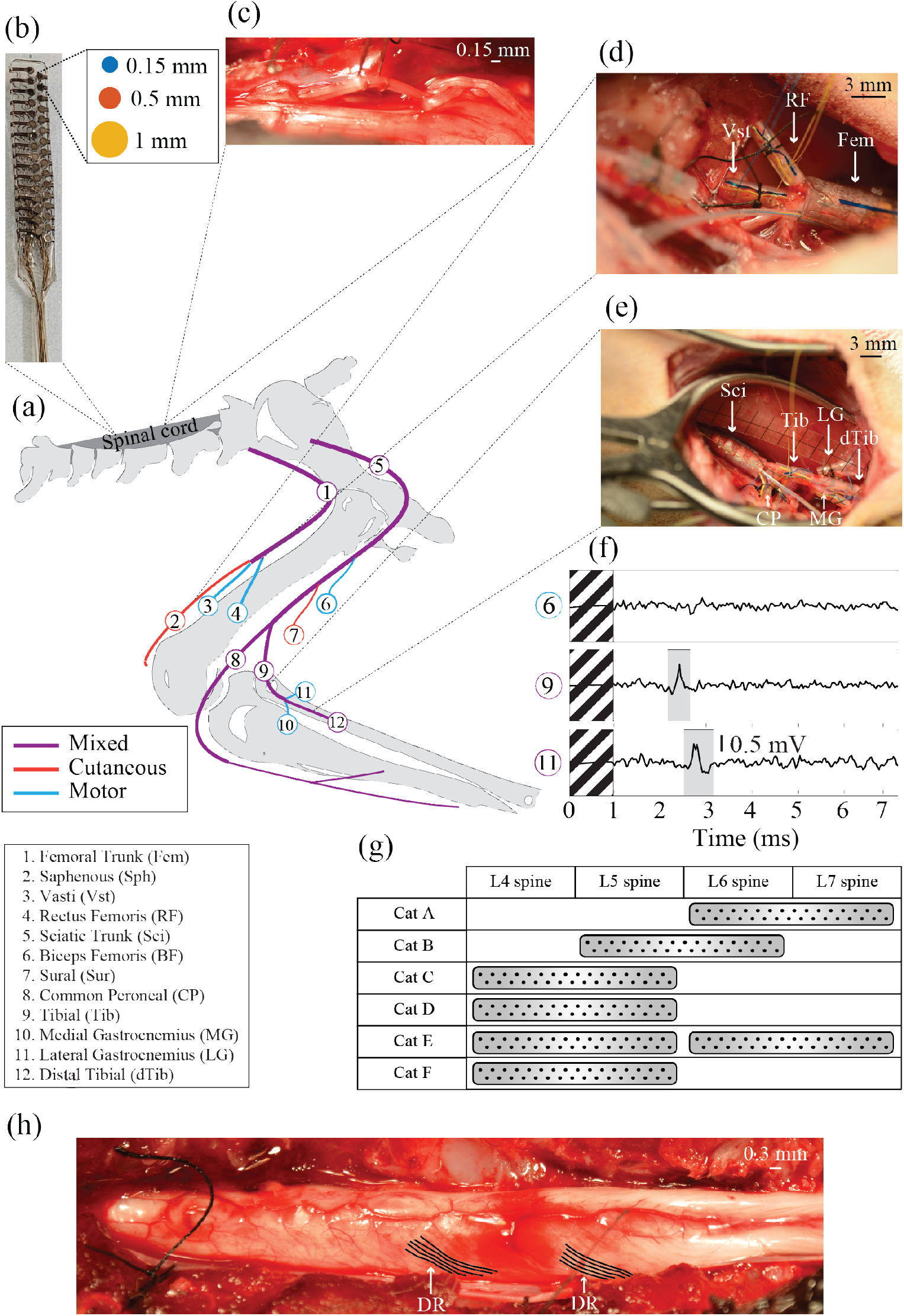
Experimental setup. (a) Schematic and list of peripheral nerves instrumented in each experiment. (b) Custom 32-channel lateral spinal cord stimulation (LSCS) paddle electrode array. Three paddle designs were used, differing only in contact diameter (0.15 mm, 0.5 mm, 1mm), as shown by the color legend. (c) Example of dorsal rootlets (DR) stimulation using bipolar hook electrodes. (d–e) Intraoperative images of nerve cuff placement on the femoral (d) and sciatic (e) branches. (f) Example CAPs recorded, with shaded window showing detection interval. (g) Summary of LSCS paddle placement across animals (Cats A–F). (h) Intraoperative dorsal view of the spinal cord after resection of the dura mater. Representative DR are indicated by arrows and manually outlined in black to improve visibility.

To verify effective contact between the nerve cuffs and the underlying nerves, we performed motor threshold after implantation of each cuff. Stimulation pulses were delivered through the cuff electrodes at gradually increasing amplitudes until visible muscle contractions were observed in the hindlimb. Once contact was confirmed, each cuff was secured in place using sutures to ensure mechanical stability during the experiment. Stainless steel disk electrodes (3-cm diameter) were placed subcutaneously over the left and right hips to serve as the recording reference and the return path for stimulation. This configuration was used consistently for both LSCS and DR stimulation.

### Lateral spinal cord stimulation

For clarity, in this study, the term spinal level refers to the neuroanatomical level of the spinal cord, defined by which spinal root projects into the cord at that location, whereas vertebral level refers to the bony vertebrae. We performed a laminectomy at vertebral levels L3–S1 to expose the lumbosacral spinal cord. During LSCS experiments, the dura mater remained intact. A custom-built 32-contact paddle electrode array (Cortec GmbH, Germany) was placed over the left lateral surface of the spinal cord (Figure 1b). Each paddle electrode consisted of a 16 × 2 array of circular platinum contacts embedded in a flexible silicone substrate that spanned approximately two vertebral levels. Contacts were arranged with a uniform vertical spacing of 2 mm and a mediolateral spacing of 1.6 mm. The array was designed to conform to the curvature of the spinal cord while maintaining consistent electrode spacing across three versions of the paddle electrode, which differed only in contact diameter (0.15, 0.5, or 1 mm). In a randomized order, each paddle was positioned over the cord and used to deliver monopolar stimulation through all 32 contacts. Stimulation was delivered using a Grapevine Neural Interface Processor (Ripple Neuro, UT) and pulses were symmetric, biphasic (66 µs/phase), and delivered while ENG signals were recorded from all instrumented peripheral nerves to characterize activation patterns. ENG recordings were performed with the same processor, using a differential headstage (Surf S2).

The vertebral level at which LSCS was performed varied across animals and determined the corresponding DR stimulation levels. A summary of electrode placements is provided in Figure 1g.

### Dorsal rootlet stimulation

Following completion of the LSCS experiments, the dura mater was carefully resected to expose the DR (Figure 1c, 1h) at the same vertebral levels previously targeted by the LSCS paddle (Figure 1g). Individual DR were elevated on a pair of hook electrodes. Bipolar stimulation was delivered through the pair of hook electrodes and CAPs were recorded from all instrumented peripheral nerves to determine which branches were activated, using the same stimulation system described above. During stimulation, DR were gently elevated using the hook electrodes to avoid contact with the phosphate-buffered saline used to maintain tissue hydration in the epidural space, minimizing current shunting and ensuring localized stimulation. Because DR are small and can often be subdivided into finer branches, we aimed to stimulate bundles that could easily be separated with minor surgical manipulation with a nerve hook. These bundles were defined as individual “rootlets” for the purpose of mapping.

### Stimulation protocols and compound action potentials detection

Custom MATLAB (R2024a) scripts were used to modulate stimulation and record CAPs in both LSCS and DR stimulation experiments. To identify responsive nerves, we used a previously established methodology [19], [20], [21]. We began each stimulation session with a high-amplitude survey, delivering 50 pulses at 20 Hz using 300–350 µA through each LSCS paddle contact or pair of DR hook electrodes. The chosen amplitude corresponded to the highest level that reliably evoked neural responses across multiple nerves without producing visible hindlimb movement. Stimulus-triggered averaging was applied to enhance the signal-to-noise ratio, and evoked CAPs were examined across all instrumented nerve branches. Only stimulation sites (contacts or hooks) that elicited a response in at least one nerve were selected for further testing. For these sites, we performed a binary search to identify the minimum stimulation amplitude required to evoke a detectable CAP. Starting with the survey amplitude, stimulation current was iteratively adjusted, and 250 pulses were delivered at each test amplitude. The ENG responses to each pulse were averaged, bandpass filtered (300 Hz– 7,500 Hz), digitized, and sampled at 30 kHz using the same recording system. To optimize stimulation frequency during the binary search, we measured the longest-latency CAP observed across all nerve cuffs, added a 5-ms buffer to prevent temporal overlap, and calculated the stimulation rate as the inverse of this value. This approach resulted in frequencies between 25 and 100 Hz across experiments, enabling efficient data collection. A binary search for all 32 electrodes on a paddle typically required 1–2 hours to complete, and a binary search during DR stimulation typically required only 3-5 minutes per rootlet.

To remove stimulation artifacts, a 0.5-ms window starting at the onset of each stimulus pulse was blanked from the recorded ENG, and the signal was interpolated between the start and end of the blanked segment. CAP detection was based on comparison to baseline (no-stimulation) ENG recordings. For each stimulation amplitude, 200 stimulation-triggered averages were computed from random subsamples comprising 80% of the 250 individual responses. A windowed root-mean-squared (RMS) amplitude was calculated for each averaged response using a 250-μs sliding window with a 25-μs step size. Baseline ENG recordings were segmented and analyzed using the same subsampling procedure to generate a distribution of baseline RMS values. The mean and standard deviation of this baseline distribution were computed across subsampling iterations. A CAP was considered present when 95% of the stimulation-triggered RMS values within a sliding window exceeded three standard deviations above the baseline mean [19]. Additionally, a subset of responses from LSCS and DR stimulation was visually inspected and validated by an experienced investigator for each animal across all instrumented nerve recordings.

### Post-processing and data analysis

All analyses and visualizations were conducted using MATLAB (R2024a) and Python Jupyter Notebooks (v7.0.8).

Threshold amplitude, defined as the minimum stimulation amplitude required to elicit a CAP in any nerve, was compared across spinal levels using the Kruskal–Wallis test. Amplitudes followed a non-normal distribution (p < 0.001, Shapiro–Wilk test), justifying the use of non-parametric methods. When significant differences were found, post hoc comparisons were performed using Dunn’s test with Bonferroni correction to account for multiple comparisons. Significance was defined as p < 0.05.

We calculated recruitment percentages for each nerve branch by determining the proportion of times a CAP was observed at or above threshold during stimulation of DR at each spinal level, across all animals. A value of 0% indicated that the nerve was never recruited at that level, while 100% indicated that it was always recruited when stimulating DR at that level.

To visualize differences in nerve recruitment during DR stimulation, responses recorded on each nerve were normalized to the maximum response amplitude observed across all DR within the same animal. This normalization enabled consistent comparisons across DR while minimizing the effect of absolute magnitude differences across nerves. Analyses were performed using responses generated at high stimulation amplitudes, which were selected to be as consistent as possible across all DR in each animal. When an identical amplitude was not available for every rootlet, the closest available amplitude near the upper end of the range of stimulation amplitudes tested in each animal was used. Additionally, because there were varying numbers of DR across spinal levels and across animals, a pooled analysis was performed across all animals by interpolating the rootlet responses for each animal to 10 evenly spaced points per level per animal. This interpolation standardized the rostrocaudal sampling of rootlet responses within each level and enabled responses from different animals to be combined for group-level comparisons.

Dynamic range analysis was performed for both DR and LSCS to evaluate the range of stimulation amplitudes over which selective nerve recruitment could be maintained. Dynamic range was calculated only when a nerve was recruited selectively (i.e., only one nerve branch was activated at threshold). Dynamic range was defined as the difference between the smallest stimulation amplitude required to recruit a single nerve branch and the amplitude at which an additional nerve branch was next recruited [15]. DR dynamic range values were computed across all cats and spinal levels in this study, whereas LSCS dynamic range values were obtained from the companion paper that describes selectivity of epidural LSCS [15] and are included here for comparison. For LSCS, dynamic range values were calculated separately for each contact size (0.15, 0.5, and 1 mm) across all cats and spinal levels. Dynamic range values followed a non-normal distribution (p < 0.001, Shapiro–Wilk test) and were compared using the Kruskal–Wallis test. When significant, post hoc comparisons were conducted using Dunn’s test with Bonferroni correction.

To assess the similarity of nerve recruitment patterns between LSCS and DR stimulation, we computed a similarity index (SI_i,j_) for each pair of paddle contacts (i) and rootlets (j). For the set of nerves recruited at threshold by LSCS (LSCS_Thresh_(i)) and DR stimulation (DR_Thresh_(j)), this index was calculated as:

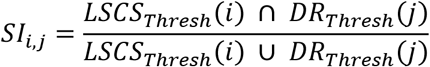

providing a measure of overlap in afferent pathways activated by each stimulation modality. The similarity index ranged from 0 to 100, where 0 indicated that the contact and rootlet recruited no common nerves, and 100 indicated complete overlap in recruited nerves.

## RESULTS

We performed neurophysiology experiments in six cats to characterize DR recruitment patterns and their relationship to LSCS activation with the goal of understanding how the functional organization of the dorsal roots affects spatial selectivity during epidural stimulation.

### Threshold amplitudes and recruitment percentages across spinal levels during dorsal rootlet stimulation

Threshold amplitudes during DR stimulation exhibited a clear rostrocaudal organization between femoral and sciatic branches, with lower thresholds observed at rostral spinal levels (L4–L5) for femoral branches and at more caudal levels (L6–L7) for sciatic branches (Figure 2a). For instance, at L4-L5, femoral branches were typically recruited at thresholds below ∼100 µA, whereas sciatic branches required higher thresholds, with recruitment observed up to ∼300 µA. This organization is consistent with LSCS recruitment patterns reported in our companion paper [15], in which stimulation at rostral vertebral levels (L4–L5) preferentially activated femoral branches and caudal levels (L6–L7) preferentially activated sciatic branches, suggesting that DR follow a similar rostrocaudal arrangement. Statistical comparisons (Kruskal–Wallis with Dunn’s post hoc test and Bonferroni correction) confirmed significant differences in thresholds across spinal levels for multiple nerves (Figure 2a).

**Figure 2:**
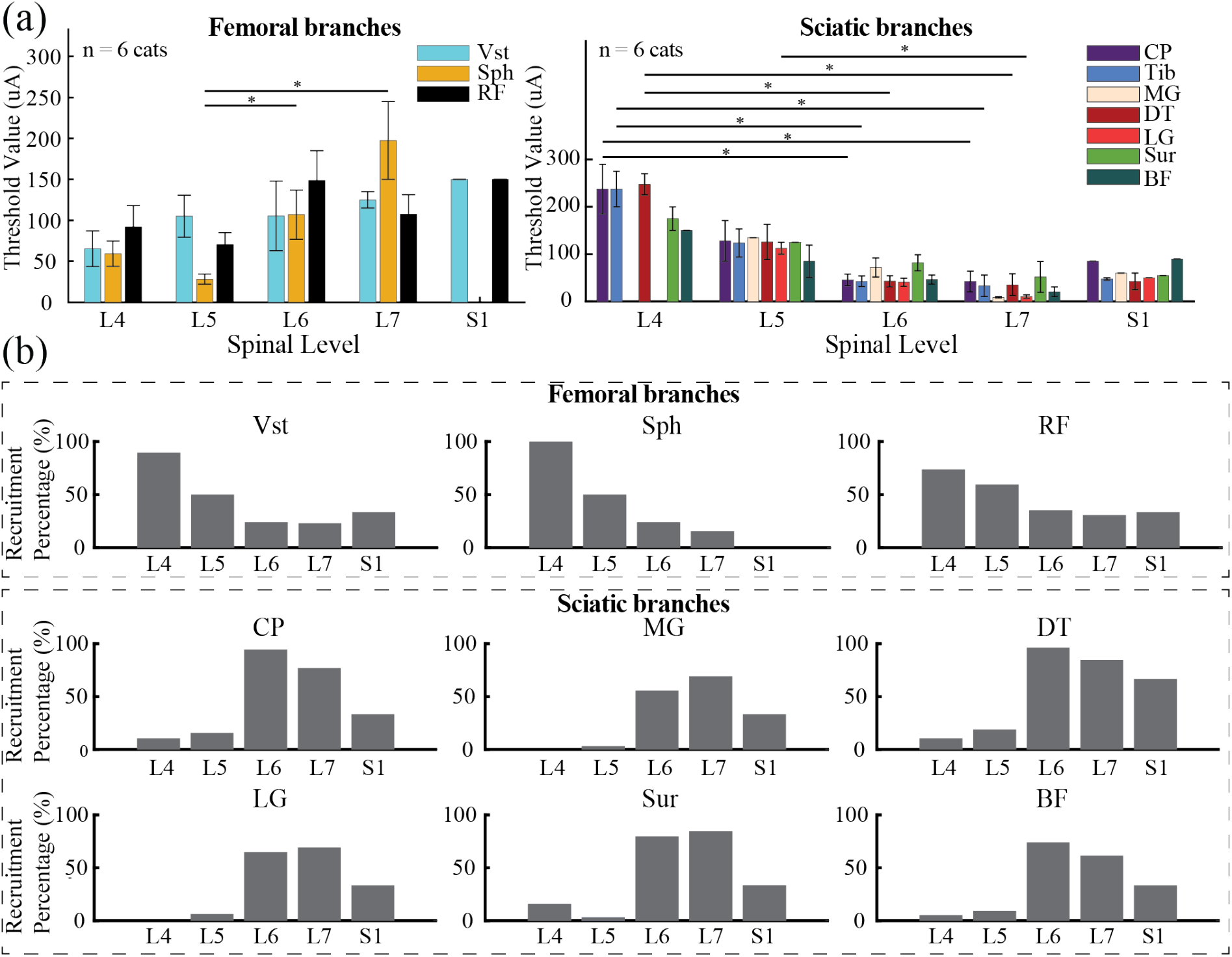
Threshold amplitudes and recruitment percentages during dorsal rootlet (DR) stimulation across spinal levels. (a) Threshold amplitudes for femoral (Vasti [Vst], Saphenous [Sph], Rectus Femoris [RF]) and sciatic (Common Peroneal [CP], Tibial [Tib], Medial Gastrocnemius [MG], Distal Tibial [DT], Lateral Gastrocnemius [LG], Sural [Sur], Biceps Femoris [BF]) nerve branches across spinal levels (n = 6 cats). Statistical comparisons were performed using the Kruskal–Wallis test, followed by Dunn’s post hoc test with Bonferroni correction for multiple comparisons (p < 0.05). Error bars represent standard error of the mean. (b) Recruitment percentages for each nerve branch at different spinal levels, expressed as the proportion of times a nerve was recruited at or above threshold across all animals. A value of 0% indicates that the nerve was never recruited at that level, while 100% indicates that it was always recruited when stimulating rootlets at that level.

Recruitment percentages further supported this rostrocaudal trend (Figure 2b). Femoral branches were recruited in ∼70–90% of stimulations at L4–L5 spinal levels but rarely at L6–L7, whereas sciatic branches showed the opposite pattern, with recruitment exceeding ∼70–80% at L6–L7. Data from S1 were limited to a single animal and were therefore excluded from cross-level comparisons.

### Nerve responses during dorsal rootlet stimulation

Although DR stimulation demonstrated a clear rostrocaudal organization between femoral and sciatic branches (Figure 2), no evidence of somatotopic organization was observed within these groups. Normalized responses from a representative animal (Cat E; Figure 3a) showed that stimulating a single rootlet typically activated multiple nerves within the same group: rostral DR recruited several femoral branches, whereas caudal DR similarly co-activated multiple sciatic branches. Importantly, these responses were recorded at high stimulation amplitudes that were comparable across DR within each cat, reducing the likelihood that differences in recruitment were due to stimulation intensity. This lack of fine spatial specificity was consistent across animals, as shown in Supplementary Figure 1.

**Figure 3:**
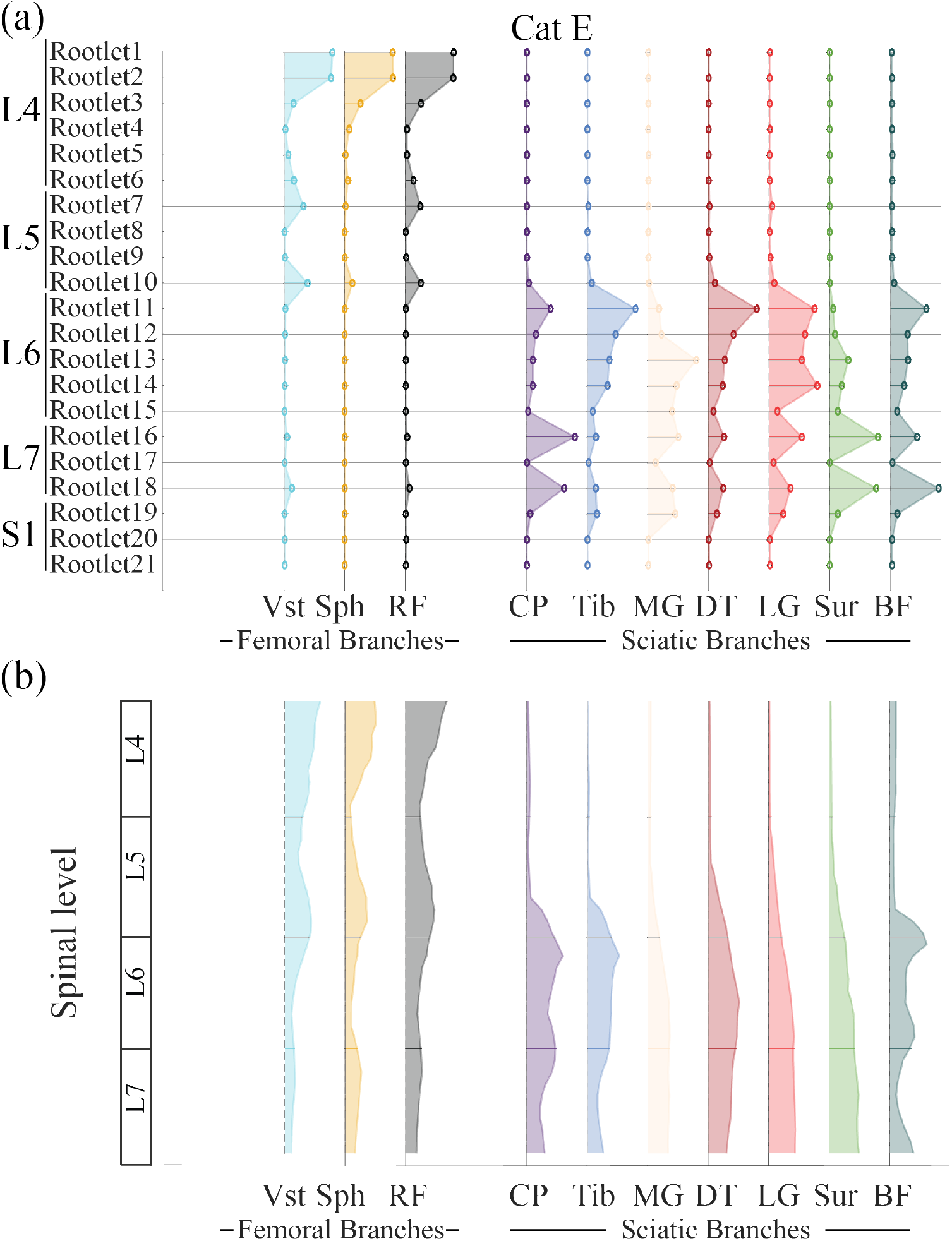
Nerve responses during dorsal rootlets (DR) stimulation. (a) Normalized nerve responses from a single cat (Cat E) while stimulating individual rootlets. By normalizing each nerve’s responses to its own maximum response amplitude across all rootlets, differences in absolute response magnitudes between nerves were accounted for, allowing comparison of relative recruitment patterns across rootlets. The highest response for each nerve was set to 1, and all other responses were expressed as a fraction of this maximum. Each column represents a single nerve branch, with femoral branches shown on the left and sciatic branches on the right. Multiple nerves within the same group were commonly co-activated by individual rootlets, with no evidence of somatotopic organization within femoral or sciatic branches. (b) Pooled analysis across all cats. Normalized responses were interpolated to 10 evenly spaced points per spinal level for each animal and combined to illustrate group-level recruitment trends. Femoral responses predominated at rostral spinal levels (L4–L5), whereas sciatic responses were more prominent at caudal levels (L6–L7). Femoral activation observed at L6–L7 likely reflects elevated noise in femoral branch recordings from Cat A, rather than genuine caudal recruitment.

A pooled analysis across all animals confirmed this pattern (Figure 3b). Normalized responses interpolated across DR at each spinal level revealed broad and overlapping recruitment of nerves within both femoral and sciatic groups. Some femoral responses appeared at caudal levels (L6–L7) in the pooled plot; however, this was likely because of elevated noise levels in Cat A’s femoral recordings (Supplementary Figure 1) rather than true caudal recruitment.

### Dynamic range during dorsal rootlet and lateral spinal cord stimulation

DR stimulation exhibited very narrow dynamic ranges, with a median of 10 µA across all cats and spinal levels (Figure 4a).

**Figure 4:**
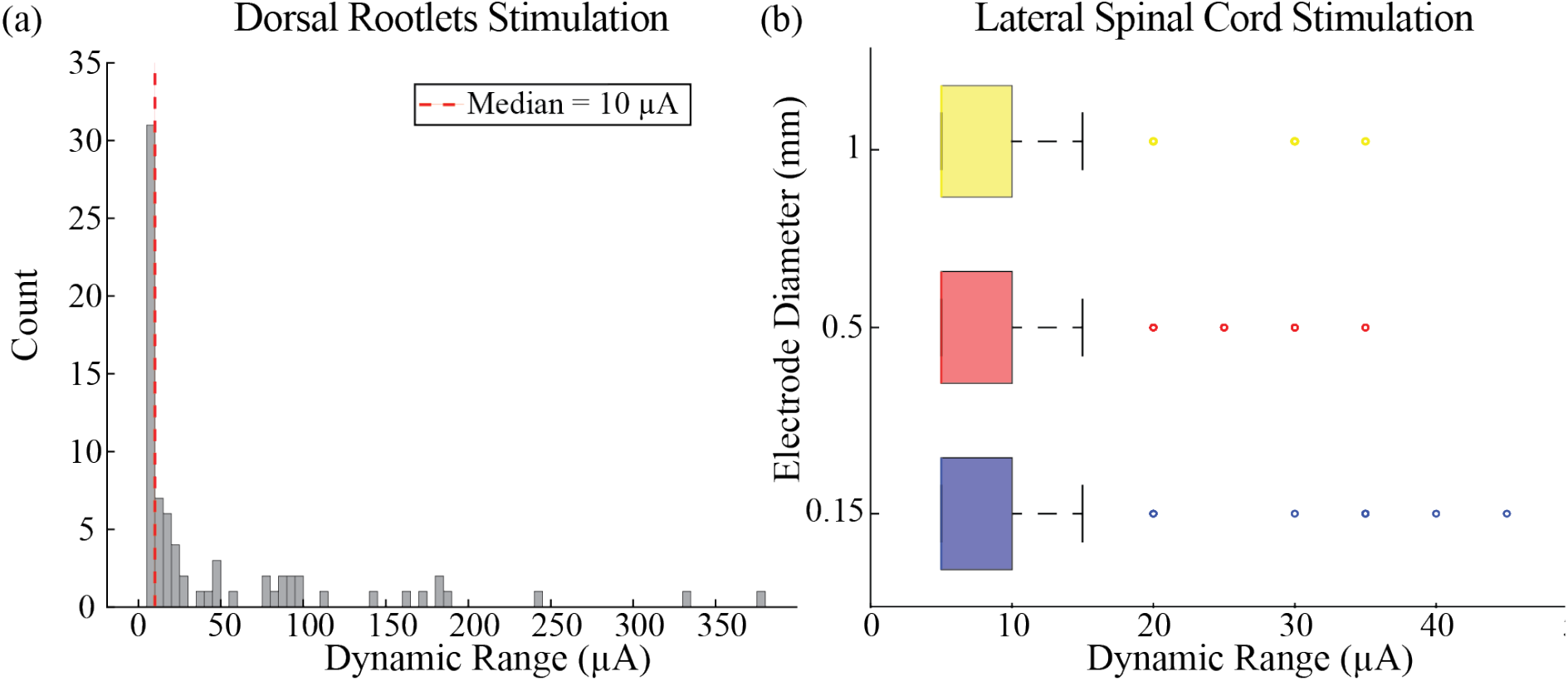
Dynamic range during dorsal rootlets (DR) and lateral spinal cord stimulation (LSCS). (a) Dynamic range distribution during DR stimulation. Histogram showing the dynamic range calculated across all cats and spinal levels. The median dynamic range was 10 µA (red dashed line), indicating that selective recruitment of a single nerve branch could only be maintained over a very narrow amplitude range before additional nerves were recruited. (b) Dynamic range during LSCS. Boxplots show dynamic range values across all contacts and spinal levels for 0.15 mm (blue), 0.5 mm (red), and 1 mm (yellow) electrode contacts. Median dynamic ranges were 5 µA for all contact sizes, with no significant differences between them (Kruskal–Wallis, p > 0.05).

Dynamic ranges were even smaller during LSCS (Figure 4b). Median dynamic range values were 5 µA for 0.15, 0.5, and 1 mm contacts, with no significant differences between contact sizes (Kruskal– Wallis, p > 0.05). However, the dynamic ranges for all LSCS configurations were significantly narrower than those observed with DR stimulation (Kruskal–Wallis followed by Dunn’s test with Bonferroni correction, all p < 0.001).

### Similarity of nerve recruitment between dorsal rootlets and lateral spinal cord stimulation

Recruitment patterns observed during LSCS closely mirrored those elicited by DR stimulation, indicating that LSCS engages similar afferent pathways at corresponding spinal levels. Recruitment curves from a representative animal (Cat C) demonstrated that nerves were recruited in a similar sequence and to comparable extents across the two stimulation modalities, despite differences in absolute thresholds (Figure 5a). As expected, DR stimulation generally required lower thresholds than LSCS because current was delivered directly to the DR. However, the overall recruitment profiles were highly comparable: at the L5 spinal level, stimulation of rootlet #6 and LSCS contact #7 (1-mm contact size) recruited the same set of femoral branches in a similar order, and at the L6 spinal level, rootlet #24 and LSCS contact #27 (1 mm) showed analogous recruitment of sciatic branches. Interestingly, when comparing regions of high similarity (white boxes in Figure 5b) to the borders of spinal levels (red lines in Figure 5b), there is a downward and rightward shift in the plots, which likely reflects the difference between vertebral and spinal levels. At the lumbar spine, spinal levels are more caudal than vertebral levels because the spinal roots project rostrally from the foramen before entering the spinal cord at the dorsal root entry zone. Since we labeled LSCS electrodes based on vertebral level and DR based on spinal level, this results in a caudal offset between DR and LSCS electrodes, corresponding to a rightward and downward shift in the plots.

**Figure 5:**
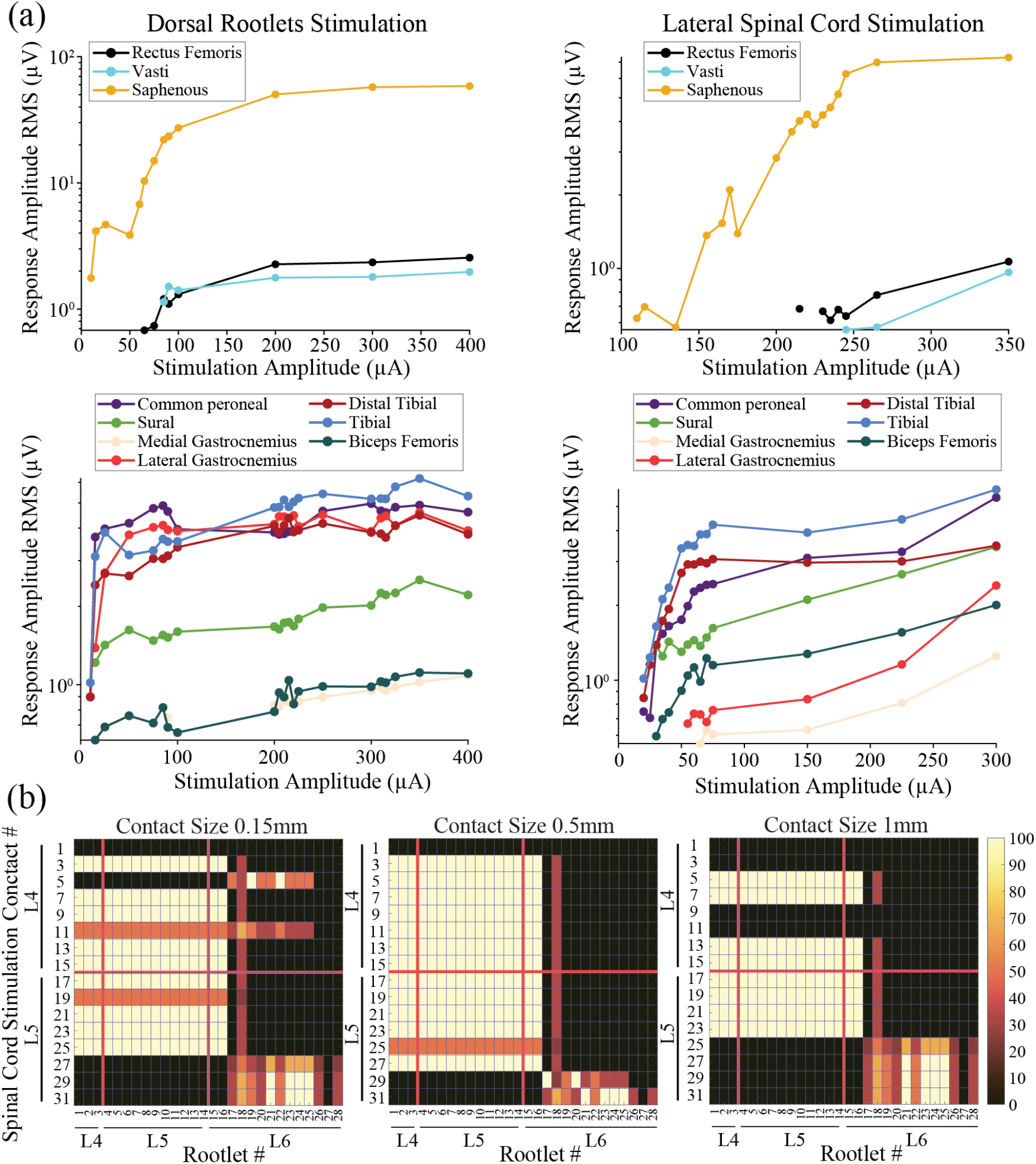
Comparison of nerve recruitment between dorsal rootlets (DR) and lateral spinal cord stimulation (LSCS). (a) Representative recruitment curves (RMS amplitude of evoked compound action potentials as a function of stimulation amplitude) from Cat C comparing DR stimulation (left) and LSCS (right) at matched spinal levels. Top: recruitment curves for rootlet #6 at the L5 spinal level and LSCS contact #7 (1-mm contact) aligned to the same vertebral region. Bottom: recruitment curves for rootlet #24 at the L6 spinal level and LSCS contact #27 (1-mm contact). Despite lower thresholds for DR stimulation, recruitment patterns were highly similar between DR and LSCS, with comparable order and extent of nerve activation. (b) Similarity index between DR stimulation and LSCS for Cat C. The index quantifies the overlap in recruited nerves at threshold for each rootlet–contact pair, expressed as the intersection over the union of recruited nerves. Values range from 0 (no shared nerves) to 100 (identical recruitment). Similarity values transitioned rostrocaudally from high femoral nerve recruitment to high sciatic nerve recruitment, reflecting the underlying DR organization.

For our analyses, we assumed that the DR were approximately equidistant and evenly distributed along the vertebral levels spanned by the LSCS paddle (L4–L5 vertebrae in this animal), so that we could pair stimulation contacts with nearby DR.

A similarity analysis further confirmed this correspondence (Figure 5b). The similarity index, a measure of overlap between peripheral nerves recruited by stimulation of each DR and through each LSCS electrode, showed high overlap at matched spinal levels, with femoral branches predominating at rostral levels and sciatic branches at caudal levels. Together with the narrow dynamic ranges reported above, these findings suggest that LSCS activates DR in a manner that largely reflects their underlying anatomical organization, indicating that LSCS selectivity is constrained by the lack of somatotopic organization within the DR.

## DISCUSSION

LSCS has shown promises for restoring sensory feedback in lower-limb amputees, but its spatial selectivity remains limited, with percepts often spreading to unintended regions in the residual limb [13], [14]. In our companion study [15], reducing electrode contact size from 2.5 to 1 mm improved selectivity, but further reduction to 0.5 mm and 0.15 mm did not result in further improvements, suggesting that anatomical factors, rather than electrode design, may constrain focal activation. To investigate these potential anatomical limitations, we directly stimulated individual DR to characterize their organization and assess how it relates to LSCS recruitment patterns.

Our experiments revealed a clear rostrocaudal organization between femoral and sciatic branches, consistent with LSCS recruitment patterns at corresponding vertebral levels [15]. However, no somatotopic organization was observed within femoral or sciatic groups, as single DR frequently co-activated multiple branches within the same group. Dynamic range analysis confirmed that selective recruitment could only be maintained over a very narrow amplitude window, and LSCS exhibited even smaller dynamic ranges than DR stimulation. Recruitment curves and similarity analysis further demonstrated that LSCS activates DR in a manner closely reflecting their anatomical organization, confirming that LSCS selectivity is constrained by the intrinsic organization of the DR rather than electrode design. The absence of somatotopic organization within femoral and sciatic branches suggests that even with highly localized stimulation (e.g., at the rootlet level), co-activation of multiple nerves is unavoidable. This intrinsic organization of the DR likely imposes fundamental limits on the focal selectivity achievable with LSCS.

Our findings are consistent with early neurophysiological mapping studies [22], which reported a serial arrangement of sensory afferents entering the dorsal roots that corresponded to major limb regions, along with shifting overlap between successive rootlet fields within a dermatome. In a similar manner, we observed a clear rostrocaudal organization between femoral and sciatic branches and overlapping recruitment within each group. Although individual DR often co-activated multiple peripheral nerve branches, both studies point to a coarse organization of the DR with limited fine-grained somatotopy. The general agreement between these independent approaches— mechanical receptive field mapping and direct electrical stimulation—supports the view that the DR are organized primarily at a gross functional level rather than by tightly segregated peripheral territories.

Additionally, the strong correspondence between DR and LSCS recruitment patterns observed in this study underscores that LSCS performance is ultimately capped by the anatomical organization of the DR. The suprathreshold DR results demonstrated broad overlapping recruitment within femoral and sciatic groups confirming that focal selectivity cannot be consistently maintained at high stimulation amplitudes. Once a single nerve branch was selectively recruited, additional branches were activated with only a slight increase in stimulation amplitude, leaving little margin to maintain selective recruitment. Despite this limitation, LSCS can still achieve selective activation of individual nerves, but only within a very narrow range of stimulation amplitudes around threshold [15]. This selectivity likely arises because, near threshold, LSCS may recruit only a subset of sensory afferents within a single rootlet, effectively exploiting sub-rootlet level selectivity before additional afferents are activated at higher currents. The narrower dynamic range observed with LSCS compared to DR stimulation indicates that LSCS cannot achieve finer selectivity than direct rootlet stimulation, further supporting the idea that focality is constrained by the anatomical organization of the DR rather than electrode design. Given this anatomy, moving the electrodes intradurally would not be expected to substantially improve focality, as stimulation would still recruit multiple overlapping nerves once suprathreshold levels are reached. These findings collectively suggest that LSCS performance is bounded by the coarse somatotopy of the DR, with improvements in percept localization more likely to result from optimized paddle placement to exploit the rostrocaudal organization rather than further reductions in contact size.

### Limitations and future work

This study has several important limitations. First, all experiments were conducted in cats, and while they are commonly used as a model for the human sensory system, species differences in DR organization may limit direct translation to humans. Second, the experiments were performed in acute preparations, and we recorded antidromic CAPs rather than functional or perceptual outcomes. Although CAPs provide an objective measure of afferent recruitment, they do not necessarily capture how these recruitment patterns translate into sensory percepts or functional improvements during LSCS. Third, our experiments were limited to lumbar DR; whether similar organizational principles hold in other spinal levels, where functional somatotopy may differ, remains to be determined.

Future work should focus on strategies that maximize the functional benefits of LSCS within the anatomical constraints of the DR. Proper paddle electrode placement to target the appropriate spinal levels—rostral for femoral afferents and caudal for sciatic—is likely to have a greater impact on percept localization than further reductions in contact size. Computational modeling that integrates rootlet-level anatomy with realistic electrical fields may help guide such placement and inform current-steering strategies. Despite these anatomical limitations, LSCS has already demonstrated meaningful functional improvements in humans [13], [14], including enhanced prosthesis control, balance, and reduced phantom limb pain. These clinical benefits underscore the relevance of LSCS, even if perfectly focal percepts cannot be consistently achieved, and motivate continued refinement of stimulation strategies that leverage the coarse but reliable rostrocaudal organization of the DR.

## ACKNOWLEDGMENTS

The authors thank Rachel Pitzer and Tyler Simpson for their assistance in preparing and performing the experiments. We also thank Cortec GmbH (Germany) for their support in designing and manufacturing the LSCS paddle and nerve cuff electrodes. This work was supported by the National Institutes of Health (NIH) (R01NS121028).

## SUPPLEMENTARY MATERIAL

**Supplementary Figure 1:**
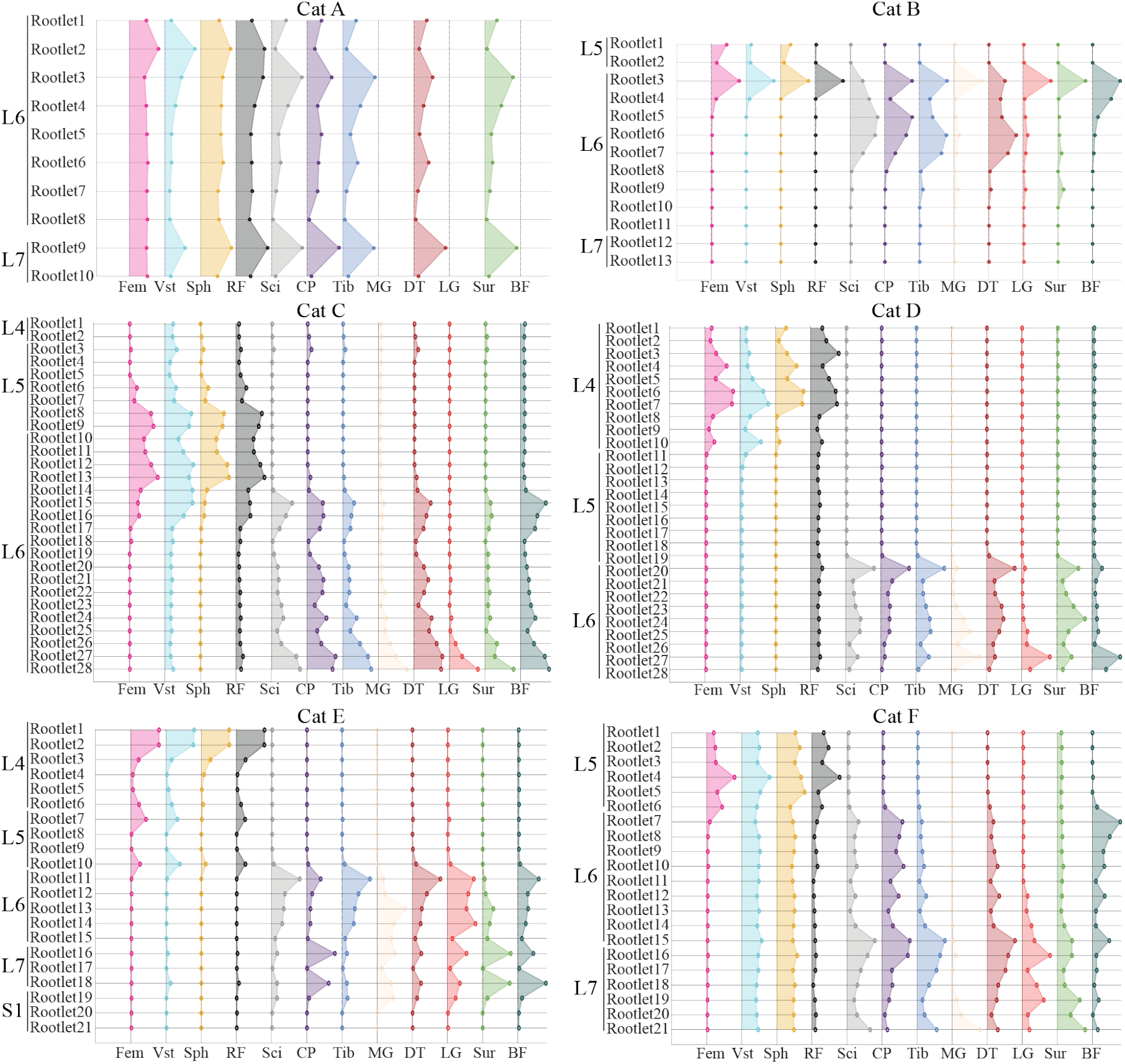
Normalized nerve recruitment patterns across individual dorsal rootlets (DR) for all six cats. Each plot shows the normalized compound action potential (CAP) amplitudes recorded from femoral (Fem), vasti (Vst), saphenous (Sph), rectus femoris (RF), sciatic (Sci), common peroneal (CP), tibial (Tib), medial gastrocnemius (MG), distal tibial (DT), lateral gastrocnemius (LG), sural (Sur), and biceps femoris (BF) nerves during DR stimulation at high stimulation amplitudes, comparable for each cat. Rootlets are arranged rostrocaudally within each spinal level (L4–S1). Broad and overlapping recruitment patterns within femoral and sciatic groups are evident across cats, demonstrating the lack of fine-grained somatotopic organization within functional groups.

